# Differential associations of various depression-related phenotypes with cardiometabolic risks: Identification of shared genetic factors and implications for drug repositioning

**DOI:** 10.1101/140590

**Authors:** Brian Chi-Fung Wong, Carlos Kwan-Long Chau, Fu-Kiu Ao, Cheuk-Hei Mo, Sze-Yung Wong, Yui-Hang Wong, Hon-Cheong So

**Author notes:** Correspondence to: Hon-Cheong So, Lo Kwee-Seong Integrated Biomedical Sciences Building, The Chinese University of Hong Kong, Shatin, Hong Kong.

## Abstract

Numerous studies have suggested associations between depression and cardiometabolic abnormalities or diseases, such as coronary artery disease and type 2 diabetes. However, little is known about the mechanism underlying this comorbidity, and whether the relationship differs by depression subtypes. Using the polygenic risk score (PRS) approach and linkage disequilibrium (LD) score regression, we investigated the genetic overlap of various depression-related phenotypes with a comprehensive panel of 20 cardiometabolic traits. GWAS results for major depressive disorder (MDD) were taken from the PGC and CONVERGE studies, with the latter focusing on severe melancholic depression. GWAS results on general depressive symptoms (DS) and neuroticism were also included. We also identified the shared genetic variants and inferred enriched pathways. In addition, we looked for drugs over-represented among the top shared genes, with an aim to finding repositioning opportunities for comorbidities.

We found significant polygenic sharing between MDD, DS and neuroticism with various cardiometabolic traits. In general, positive polygenic associations with CV risks were observed for most depression phenotypes except MDD-CONVERGE. Counterintuitively, PRS representing severe melancholic depression was associated with reduced CV risks. Enrichment analyses of shared SNPs revealed many interesting pathways, such as those related to inflammation, that underlie the comorbidity of depressive and cardiometabolic traits. Using a gene-set analysis approach, we also revealed a number of repositioning candidates, some of which were supported by prior studies, such as bupropion and glutathione. Our study highlights shared genetic bases of depression with cardiometabolic traits, and suggests the associations vary by depression subtypes. To our knowledge, this is the also first study to make use of human genomic data to guide drug discovery or repositioning for comorbid disorders.

## Introduction

Major depression is a common psychiatric disorder worldwide. More than 300 million people globally suffer from major depressive disorder (MDD)^1^ and it is ranked as the leading cause of disability worldwide by the World Health Organization in a latest report^1^. Depression has been shown to be associated with numerous physical illnesses, including cardiovascular and metabolic diseases^2^, as well as other cardiovascular (CV) risk factors such as obesity^3^ and dyslipidemia^4^. A book-length review on this subject was given in Baune and Tully^5^. In addition, studies have also suggested a link between neuroticism and poor cardiovascular outcomes^6,7^. However, the mechanism underlying the relationship between depression and cardiometabolic traits is not well understood. As common biological pathways may underlie both depression and cardiometabolic traits^5^, shared genetic factors may possibly contribute to the comorbidities. Studies have shown genetic overlap between depression and blood pressure, total cholesterol level, body mass index (BMI), as well as heart rate variability ^8-11^. However, only a limited spectrum of cardiometabolic abnormalities has been considered in previous studies. These studies were mainly twin- or family-based and the main aim was to investigate an overall shared genetic basis; as such, they did not identify which genetic variants are most likely shared or the pathways involved. A recent systematic review nicely summarizes the current evidence for susceptibility genes shared between depression and cardiometabolic traits from candidate gene and some genome-wide studies^12^. Nevertheless, the review mainly focused on a qualitative assessment of the evidence and the results of candidate gene studies, which comprised a substantial part of the review, may not as reliable as large-scale GWAS^13^.

Depression is widely regarded as a heterogeneous condition^14^. Depression has a lifetime prevalence of up to 17% according to a US study^15^ and the diagnosis is entirely based on clinical symptoms with no reliable biomarkers. As such, it is likely that substantial genetic and phenotypic heterogeneity underlie this diagnosis. A meta-analysis on the association of depression with cardiovascular disease risks also remarked on the high level of heterogeneity among studies^16^, which may be partially attributed to the heterogeneity of depression itself. A recent twins study on the genetic overlap of depression with type 2 diabetes mellitus (DM) suggested that the genetic factors underlying the comorbidity might be different in males and females^17^. Despite the well-known heterogeneity of depression, few epidemiological studies have investigated the cardiovascular risks of different depression subtypes (or different phenotypes related to depression). Lamers et al.^18^ studied a Dutch sample of depression cases and controls, and found that patients with atypical depression had worse metabolic profiles than controls or those with melancholic depression. Lasserre et al.^19,20^ reported that only the atypical MDD subtype was prospectively associated with a raised incidence of metabolic syndrome as well as greater increase of fasting glucose and waist circumference during follow-up. However, to our knowledge, no previous studies have compared the *genetic overlap* between different depression-related phenotypes or disease subtypes with CV risk factors; in addition, the individual genes and pathways contributing to the possible genetic overlap remain to be explored. Previous epidemiological studies are also often limited by modest sample sizes (for example, the number of melancholic cases was 111 in Lamers et al^18^. and 369 in Lasserre et al.^20^).

In this study we made use of four GWAS datasets of depression or depression-related phenotypes and investigated the genetic overlap with a comprehensive panel of 20 cardiometabolic traits. These four datasets include: (1) the MDD-CONVERGE study^21^, a relatively homogenous group of Chinese women with severe and mostly (∼85%) melancholic depression; (2) the MDD-PGC study^22^ with no enrichment of particular disease subtypes; (3) a GWAS meta-analysis on depressive symptoms conducted by the Social Science Genetic Association Consortium (SSGAC-DS)^23^ which include (i) case-control (i.e. clinical depression vs no depression); and (ii) population-based samples (measuring general depressive symptoms instead of clinical diagnosis); (4) a GWAS meta-analysis by SSGAC on neuroticism^23^, a personality trait with established associations with depression^24,25^. We wish to test whether different depression-related phenotypes would have similar or diverse patterns of associations with cardiometabolic traits.

Our scheme of analyses is summarized below. Firstly, we studied whether polygenic risk score (PRS) of depression phenotypes are associated with a comprehensive panel of 20 cardiometabolic traits and vice versa; as another approach to assess genetic overlap, we also applied cross-trait LD score regression. Secondly, we identified the genetic variants most likely shared between the disorders, by a statistical approach based on local true discovery rates. Based on the shared genetic markers, we also inferred the enriched biological pathways and gene ontology terms, and drugs potentially linked to both kinds of disorders. To our knowledge, this is the also first study to make use of human genomic data to guide drug discovery or repositioning for comorbid disorders.

## METHODS

### Study samples

We studied polygenic sharing of various depression phenotypes (MDD-CONVERGE, MDD-PGC, depressive symptoms, neuroticism) with a panel of cardiometabolic traits. The four depression phenotypes have been described above. Note that the GWAS summary results on depressive symptoms also contains MDD-PGC as part of the sample, but the rest (∼90%) of the sample originated from other cohorts. Cardiometabolic traits under study included body mass index (BMI)^26^, waist-hip ratio (WHR) ^27^, fasting glucose (FG)^28^, fasting insulin (INS)^28^, insulin resistance (HOMA-IR)^29^, fat-percentage ^30^, high-density lipoprotein (HDL), low-density lipoprotein (LDL), triglycerides (TG), total cholesterol (TC) ^31^, adiponectin^32^, leptin^30^, systolic blood pressure (SBP)^33^, diastolic blood pressure (SBP)^33^, coronary artery disease (CAD) ^34^ and type 2 diabetes mellitus (DM)^35^.

GWAS summary statistics were downloaded from the Psychiatric Genomics Consortium website (<u>https://www.med.unc.edu/pgc</u>) and LD hub (<u>http://ldsc.broadinstitute.org/</u>). Details of each study may be found in the references listed above. For SBP and DBP, we used data from a meta-analysis of exome-sequencing studies^33^ as we were unable to obtain publicly available GWAS data with effect size directions.

### Polygenic score analysis

Polygenic risk score (PRS) is a weighted sum of allelic counts, with the weights given by log odds ratios of individual SNPs. The PRS for an individual *i,* or *r*_i_, can be computed by 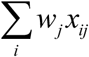, where *x*_*ij*_ is the centered allelic count for the *j*th SNP for the *i*th individual, *w*_*j*_ is the weight given to the *j*th SNP, given by the log odds ratio or regression coefficient in a linear regression. In this study we performed analyses using the summary statistics of each pair of traits, as raw genotype data are not available. Association testing was carried out by the method “gtx” described by Johnson^36^. This method was also described and elaborated in a number of papers e.g. ref.^37-39^. Briefly, we tested for the association of the PRS derived from the first trait with a second trait *y*_*i*_ (with values centered) by regression:

*y*_*i*_*=αr*_*i*_

Then it can be shown that the regression coefficient *α* can be estimated by

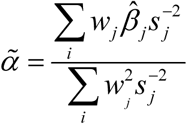

and the standard error cam be estimated by

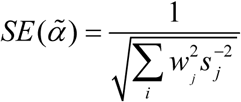

where 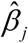 is the regression coefficient when the second trait *y*_*i*_ is regressed onto the *j*th SNP and *s*_*j*_ is corresponding standard error of *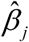*. This method was implemented in the R program PRsice^40^. We performed LD-clumping with an R^2^ threshold of 0.05 prior to association analyses, following the suggestions given in the PRsice vignette. A series of ten *p*-value thresholds (0.001, 0.005, 0.01, 0.03, 0.05, 0.1, 0.2, 0.3, 0.4, 0.5) was considered. Multiple testing was corrected by the false discovery rate (FDR) approach, which controls the expected proportion of false positives among the results declared to be significant. FDR-adjusted *p*-value (*q*-value) was calculated by the R program p.adjust using the Benjamini-Hochberg procedure^41^. Results with *q*-value below 0.05 are regarded as significant, and those with *q*-value below 0.1 are regarded as suggestive associations. FDR correction was performed for each pair of metabolic trait and depression phenotype separately, and as proven by Efron^42^, separate false discovery rate control for different groups of hypotheses at a level *q* still enables the overall FDR to be maintained at the level *q*.

#### Accounting for sample overlap

Polygenic score analyses are usually based on independent training and testing samples. We employed two approaches to guard against false positive associations due to potential sample overlap. The first is by checking for overlapping cohort samples, and the second is by examining the regression intercepts of cross-trait LD score regression.

It is worth noting in this context that a recent study by Burgess et al.^43^ addressed the issue of participant overlap in two-sample Mendelian randomization (MR), which employs a very similar statistical formulation to the above PRS analysis. However, in this study we focused on polygenic associations instead of strict causality, and relaxed some assumptions required by MR: pleiotropic effects were allowed and genetic variants need not be unequivocally associated with the risk factor. Burgess et al.^43^ showed that in a case-controls setting, if the risk factor (e.g. lipid level) is only measured in controls, then unbiased estimates can be obtained even in a one-sample setting (*i.e.* the risk factor measured in *all* control subjects). In addition, it was shown the bias is a linear function of the proportion of overlap; therefore, even if overlap is present but the extent is minor, the results are unlikely to be significantly affected.

In the case of MDD-PGC^22^, we checked that *none* of the quantitative cardiometabolic traits was measured in depressive patients. Participant overlap in the control subjects (in our case it is far less than a complete overlap) is unlikely to result in biased estimates. For CAD and DM, these are case-control studies and we found possible overlap with KORA and PopGen among the control samples. However, only *subsets* of these samples were included in the MDD-PGC and in CAD/DM GWAS, and whether there is genuine overlap (or the extent of overlap) is unknown. We evaluated the degree of overlap by considering the cross-trait LD score regression (LDSR) intercept^44^ (details of LDSR is given in the next section). For samples with no overlap, this intercept is usually close to zero^44^. We assessed whether the intercept is significantly different from zero by a Wald test (i.e. computing z-scores by the intercept estimate divided by its standard error).

Similarly, for neuroticism GWAS, there is possible overlap with GWAS samples of cardiometabolic traits in several cohorts (e.g. BLSA, EGCUT, ORCADES); however only subsamples of these cohorts were included for neuroticism or cardiometabolic measures, so the actual degree of overlap is unknown. Again we assessed the degree of overlap by checking cross-trait LDSR intercepts.

For MDD-CONVERGE, the sample was collected from China and there is no overlap with any other cardiometabolic traits, which were mainly measured in Caucasians. As for the GWAS meta-analyses of depressive symptoms by SSGAC, we did not find overlap of this sample with other metabolic traits; however since MDD-PGC is part of this sample, there may be a potential (minor) overlap of controls with CAD and DM, which is checked by LDSR.

As we shall see in the results section, the extent of sample overlap for most traits was *not* significant as evaluated by LDSR and hence our conclusions are unlikely to be substantially affected by overlap issues.

### LD score regression

LD score regression is another widely used approach to assess genetic correlation between complex diseases^44^. The approach is able to accommodate sample overlap and LD between variants. To assess genetic correlation of MDD-CONVERGE with other traits, we employed “popcorn”, a recently developed program which allows estimation of genetic correlation across different ethnicities^45^. LD scores across ancestries were computed from the EUR and EAS populations from the 1000 Genome Project^46^. Default settings were used for our analyses. We did not perform LD score regression for SBP and DBP studies as they were exome-sequencing studies and the LD pattern of genetic variants differs from GWAS results.

### Discovery of genetic variants associated to both MDD and cardiometabolic traits

The SNPs shared by MDD and cardiometabolic traits were identified by an approach based on the concept of local false discovery rates^47^. For each SNP, the probability of being associated with *both* traits (denoted by tdr_11_) was calculated based on the observed z-statistics. The approach is closely related to the conditional false discovery rate^48^ but here we focused on the chance of *shared* associations instead of probability of association conditioned on the other trait. We followed the same statistical formulation proposed by Chung et al.^49^, although we worked with the z-statistics instead of *p*-values. Briefly, a four-group mixture model of *z*-statistics was assumed:

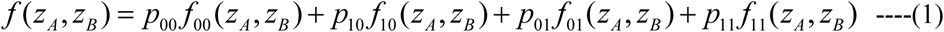

where *z*_A_ and *z*_B_ refer to the *z*-statistics of trait A and trait B respectively. There are four possibilities for each SNP when two traits are considered: 1) the SNP is associated with none of the traits; (2) the SNP is associated with the first trait only; (3) the SNP is associated with the second trait only; and (4) the SNP is associated with both traits. Each of these four possibilities are represented in the above mixture model. For instance, *p*_11_ denotes the proportion of markers that has association with both traits, and *f*_11_ denotes the probability density function of the z-statistics of this group of markers. From equation (1) we can derive the probability that a SNP is associated with both traits (denoted by *tdr*_11_), using an expectation-maximization (EM) algorithm. Further details of the method are given in Supplementary Information and in Chung et al.^49^.

### Pathway and Gene Ontology (GO) enrichment analysis by ConsensusPathDB

Shared SNPs with tdr_11_ > 0.5 (*i.e.* greater than 50% probability that the SNP is associated with *both* traits) were selected and mapped to related genes using the Bioconductor package biomaRt^50^. However, some SNPs were mapped to multiple genes, which may lead to the enrichment of certain pathways that consist of numerous genes from the same (or few) SNP(s). We adopted the following method to correct for this potential bias. We first extracted variant consequences (using sequence ontology [SO] terms) of variants via BioMart, and each SNP was mapped to the gene corresponding to the highest impact rating (as listed in <u>http://asia.ensembl.org/info/genome/variation/predicted_data.html#consequences</u>). ConsensusPathDB was used to infer the biological pathways and gene ontology (GO) (level 5) terms enriched among the top shared genes. ConsesusPathDB is a comprehensive resource that integrates multiple databases for pathway analyses^51^. Over-representation of pathways or GO terms was assessed by hypergeometric tests with a *p*-value cutoff of 0.01 with at least two genes in each pathway, following the default settings. We performed GO and pathway analysis separately for SNPs with the same and opposite directions of effects for each pair of traits; we also performed an overall analysis on all SNPs regardless of effect directions.

### Drug repositioning by over-representation analyses in WebGestalt

We then looked for drug-related gene-sets that are over-represented among the most significantly shared genes. We used WebGestalt^52^ for this analysis. WebGestalt employed a computational approach known as GLAD4U^53^ to query the scientific literature to retrieve and prioritize gene lists associated with drugs. The over-represented drugs were identified by hypergeometric tests. We require at least 2 genes in each gene-set and results with *q*-value < 0.1 were retrieved.

We expected to discover drugs associated with the overlapping genes, and therefore identify existing drugs that have genetic associations to non-target disorders. This may provide us hints for new indications of known drugs; for example, drugs known to treat psychiatric disorder can be used as a therapy for cardiometabolic diseases, or vice versa. We may also find novel repositioning candidates with beneficial effects on both depressive and cardiometabolic disorders.

## Results

### Polygenic associations between depression-related phenotypes and cardiometabolic traits

Polygenic associations of MDD-PGC with cardiometabolic traits are shown in Table 1. Firstly, PRS were constructed from the two MDD datasets, and metabolic traits are regarded as target (i.e. dependent) phenotypes. Higher polygenic score of MDD-PGC was significantly associated with lower levels of adiponectin and higher levels of TG and WHR (with or without adjustment for BMI). It was also associated with increased risks of CAD and DM at an FDR threshold of 0.05. When cardiometabolic traits were used to construct PRS and MDD-PGC was regarded as the outcome, we found associations with most of traits mentioned above, but also observed a positive relationship with HDL and inverse relationship with FG and BMI-adjusted leptin.

**Table 1.**
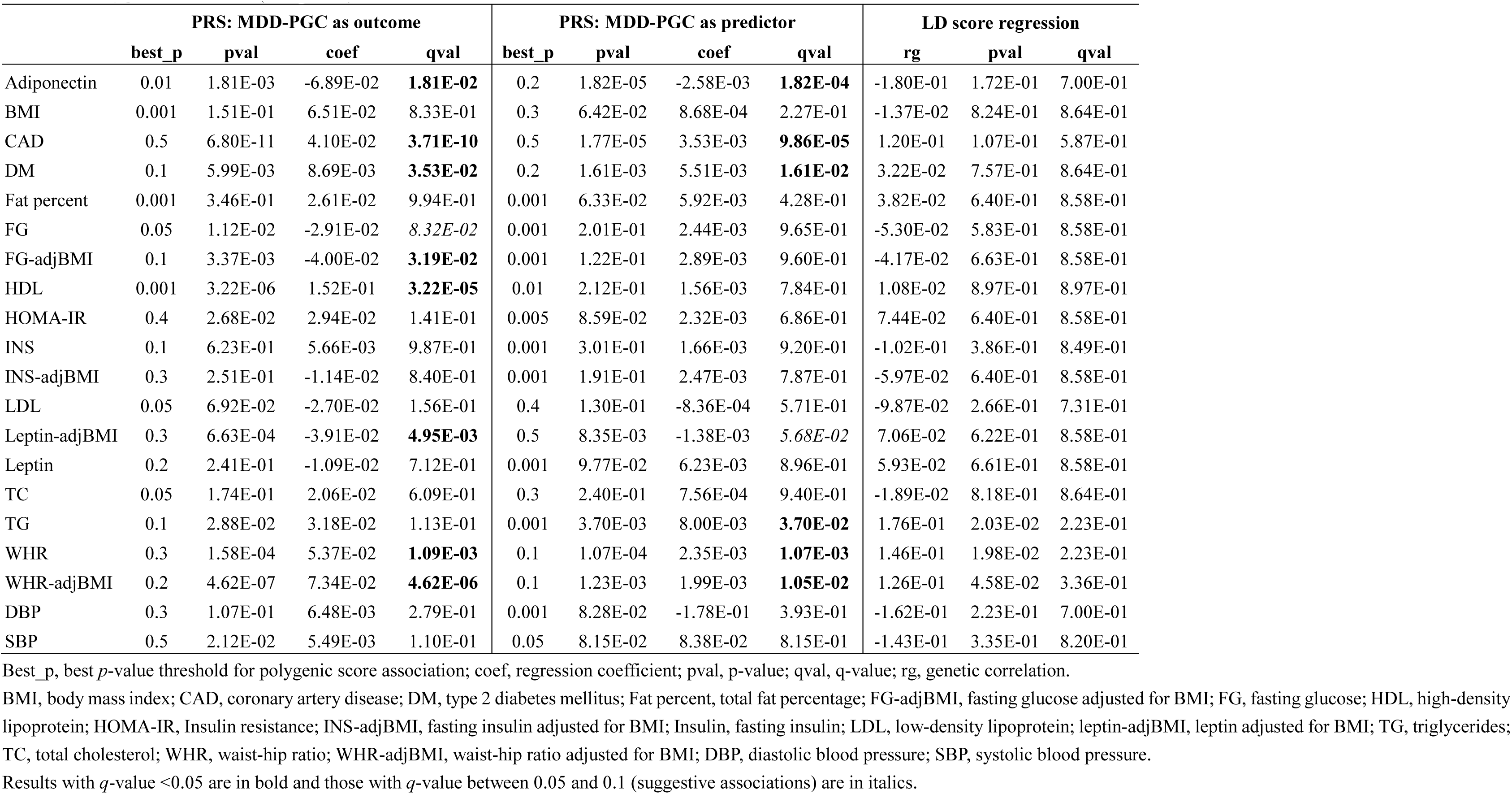
Polygenic risk score (PRS) analysis between MDD-PGC and cardiometabolic traits

As for the other GWAS study on severe melancholic depression (MDD-CONVERGE) (Table 2), surprisingly, we observed that *higher* PRS derived from MDD-CONVERGE was associated with *lower* BMI, TG, WHR (with or without adjustment for BMI), fat percentage, as well as risks of CAD and DM (with *q-*value just above 0.05 for DM); it was also associated with higher HDL. When PRS was constructed from cardiometabolic traits, we observed inverse associations with BMI, FG, TG and WHR, and positive association with HDL.

**Table 2.** Polygenic risk score (PRS) analysis between MDD-CONVERGE and cardiometabolic traits

|  | MDD-CONVERGE as outcome |  |  |  | MDD-CONVERGE as predictor |  |  |  | LD score regression |  |  |
| --- | --- | --- | --- | --- | --- | --- | --- | --- | --- | --- | --- |
|  | best_p | pval | coef | qval | best_p | pval | coef | qval | rg | p | qval |
| Adiponectin | 0.01 | 4.80E-02 | 2.24E-02 | 3.07E-01 | 0.2 | 1.41E-01 | -6.09E-04 | 4.54E-01 | 2.86E-01 | 5.72E-02 | 1.14E-01 |
| BMI | 0.03 | 1.64E-07 | -1.42E-01 | **1.19E-06** | 0.05 | 5.87E-05 | -1.72E-03 | **5.87E-04** | -1.83E-01 | 6.30E-03 | **4.49E-02** |
| CAD | 0.4 | 3.88E-02 | -1.31E-02 | 1.65E-01 | 0.5 | 5.98E-05 | -2.89E-03 | **2.28E-04** | -4.59E-02 | 4.92E-01 | 5.42E-01 |
| DM | 0.05 | 1.65E-01 | 4.25E-03 | 3.98E-01 | 0.03 | 5.18E-03 | -5.70E-03 | *5.18E-02* | -2.32E-01 | 8.90E-03 | **4.49E-02** |
| Fat percent | 0.001 | 6.59E-02 | -3.06E-02 | 4.01E-01 | 0.3 | 2.27E-03 | -1.50E-03 | **7.77E-03** | -1.85E-01 | 2.72E-02 | *8.55E-02* |
| FG | 0.3 | 2.77E-02 | -1.67E-02 | **5.58E-02** | 0.01 | 3.67E-02 | -1.14E-03 | 3.67E-01 | -2.15E-01 | 2.03E-02 | *7.44E-02* |
| FG-adjBMI | 0.01 | 1.06E-01 | -1.90E-02 | 5.03E-01 | 0.01 | 1.29E-01 | -8.49E-04 | 9.69E-01 | -1.55E-01 | 8.87E-02 | 1.63E-01 |
| HDL | 0.4 | 4.79E-05 | 4.91E-02 | **3.91E-04** | 0.5 | 4.29E-05 | 1.69E-03 | **2.97E-04** | 6.56E-02 | 4.48E-01 | 5.42E-01 |
| HOMA-IR | 0.2 | 6.96E-02 | 2.52E-02 | 3.89E-01 | 0.01 | 3.34E-02 | -1.49E-03 | 2.25E-01 | -2.08E-01 | 1.87E-01 | 2.74E-01 |
| INS | 0.005 | 2.68E-01 | -2.02E-02 | 6.77E-01 | 0.01 | 5.80E-02 | -1.05E-03 | 2.60E-01 | -1.06E-01 | 3.41E-01 | 4.41E-01 |
| INS-adjBMI | 0.1 | 1.78E-01 | 1.16E-02 | 6.48E-01 | 0.005 | 3.68E-01 | 5.23E-04 | 9.70E-01 | 2.19E-02 | 8.39E-01 | 8.79E-01 |
| LDL | 0.001 | 5.27E-01 | -2.35E-02 | 9.83E-01 | 0.001 | 1.10E-01 | 2.93E-03 | 5.62E-01 | -6.30E-03 | 9.43E-01 | 9.43E-01 |
| Leptin-adjBMI | 0.005 | 9.00E-03 | -7.44E-02 | *9.00E-02* | 0.001 | 8.65E-02 | -3.05E-03 | 5.37E-01 | -1.78E-01 | 1.72E-01 | 2.71E-01 |
| Leptin | 0.05 | 1.67E-02 | -2.78E-02 | 1.20E-01 | 0.01 | 1.99E-02 | -2.59E-03 | 1.14E-01 | -3.74E-01 | 9.30E-03 | **4.49E-02** |
| TC | 0.01 | 1.49E-01 | 3.47E-02 | 5.94E-01 | 0.001 | 8.10E-02 | 3.11E-03 | 7.63E-01 | -4.97E-02 | 4.90E-01 | 5.42E-01 |
| TG | 0.4 | 9.19E-04 | -4.07E-02 | **4.86E-03** | 0.3 | 6.77E-05 | -1.64E-03 | **3.44E-04** | -1.35E-01 | 1.02E-01 | 1.72E-01 |
| WHR | 0.01 | 3.38E-05 | -1.26E-01 | **3.38E-04** | 0.5 | 8.59E-07 | -1.79E-03 | **5.92E-06** | -1.69E-01 | 6.90E-03 | **4.49E-02** |
| WHR-adjBMI | 0.01 | 6.65E-03 | -8.33E-02 | *6.65E-02* | 0.5 | 2.02E-04 | -1.37E-03 | **1.23E-03** | -7.84E-02 | 2.12E-01 | 2.91E-01 |
| DBP | 0.01 | 3.62E-01 | 6.95E-03 | 9.97E-01 | 0.4 | 3.19E-01 | -1.22E-02 | 7.37E-01 | -2.84E-01 | 4.47E-02 | 1.14E-01 |
| SBP | 0.05 | 3.09E-01 | 2.50E-03 | 7.85E-01 | 0.1 | 1.00E-01 | -4.22E-02 | 4.78E-01 | -3.10E-01 | 1.02E-02 | **4.49E-02** |

Please refer to the legends of Table 1.

Table 3 shows the polygenic associations of depressive symptoms (from SSGAC) with cardiometabolic traits. When PRS of depressive symptoms is used as the predictor, we observed positive associations with BMI, CAD, DM, fat percentage, HOMA-IR, INS, leptin, TC, TG, WHR, SBP and DBP (Table 2). Similar associations were observed when PRS of CV risk factors were used as predictors.

**Table 3.** Polygenic risk score (PRS) analysis between depressive symptoms (DS) and cardiometabolic traits

|  | PRS: DS as outcome |  |  |  | PRS: DS as predictor |  |  |  | LD score regression |  |  |
| --- | --- | --- | --- | --- | --- | --- | --- | --- | --- | --- | --- |
|  | best_p | pval | coef | qval | best_p | pval | coef | qval | rg | pval | qval |
| Adiponectin | 0.2 | 2.70E-01 | 9.49E-04 | 7.79E-01 | 0.03 | 2.80E-02 | 9.85E-03 | 1.12E-01 | -6.90E-02 | 3.59E-01 | 4.64E-01 |
| BMI | 0.03 | 3.25E-14 | 2.65E-02 | **3.25E-13** | 0.2 | 5.77E-17 | 2.30E-02 | **3.95E-16** | 9.61E-02 | 2.27E-02 | *7.13E-02* |
| CAD | 0.1 | 1.09E-07 | 4.22E-03 | **7.36E-07** | 0.1 | 6.67E-13 | 4.04E-02 | **6.32E-12** | 1.42E-01 | 1.10E-03 | **8.07E-03** |
| DM | 0.3 | 1.02E-01 | 3.50E-04 | 3.87E-01 | 0.2 | 6.00E-08 | 5.92E-02 | **4.25E-07** | 7.03E-02 | 2.69E-01 | 4.22E-01 |
| Fat percent | 0.1 | 3.29E-04 | 2.56E-03 | **3.29E-03** | 0.2 | 2.89E-11 | 2.68E-02 | **1.17E-10** | 2.02E-01 | 2.00E-04 | **2.20E-03** |
| FG | 0.001 | 1.35E-01 | -1.04E-02 | 5.79E-01 | 0.001 | 3.62E-02 | 1.68E-02 | 3.62E-01 | 4.41E-02 | 4.31E-01 | 4.99E-01 |
| FG-adjBMI | 0.001 | 6.23E-02 | -1.24E-02 | 5.89E-01 | 0.01 | 2.09E-02 | 9.26E-03 | 1.11E-01 | 5.45E-02 | 3.51E-01 | 4.64E-01 |
| HDL | 0.4 | 1.32E-08 | -9.72E-03 | **4.93E-08** | 0.5 | 1.06E-09 | -2.02E-02 | **5.11E-09** | -9.96E-02 | 4.67E-02 | 1.28E-01 |
| HOMA-IR | 0.3 | 1.31E-05 | 8.08E-03 | **5.89E-05** | 0.2 | 3.36E-08 | 1.67E-02 | **2.26E-07** | 1.42E-01 | 1.27E-01 | 2.53E-01 |
| INS | 0.001 | 1.90E-01 | 9.65E-03 | 9.32E-01 | 0.001 | 1.11E-03 | 2.20E-02 | **1.11E-02** | 5.39E-02 | 4.87E-01 | 5.35E-01 |
| INS-adjBMI | 0.001 | 1.07E-01 | 1.07E-02 | 4.27E-01 | 0.4 | 1.56E-07 | 1.18E-02 | **6.91E-07** | 1.40E-01 | 5.22E-02 | 1.28E-01 |
| LDL | 0.5 | 1.13E-03 | 5.06E-03 | **5.65E-03** | 0.2 | 1.61E-02 | 9.16E-03 | *8.08E-02* | -7.20E-03 | 8.93E-01 | 9.35E-01 |
| Leptin-adjBMI | 0.005 | 2.05E-02 | -9.09E-03 | 2.05E-01 | 0.05 | 3.63E-01 | -4.07E-03 | 9.90E-01 | 2.23E-01 | 3.70E-03 | **1.63E-02** |
| Leptin | 0.01 | 1.21E-02 | 6.24E-03 | 1.21E-01 | 0.3 | 4.00E-03 | 1.33E-02 | **2.55E-02** | 8.95E-02 | 2.51E-01 | 4.22E-01 |
| TC | 0.3 | 4.84E-05 | 6.60E-03 | **4.84E-04** | 0.2 | 1.03E-03 | 1.22E-02 | **5.48E-03** | 3.70E-03 | 9.35E-01 | 9.35E-01 |
| TG | 0.001 | 1.63E-06 | 2.71E-02 | **1.17E-05** | 0.1 | 2.35E-11 | 2.48E-02 | **2.35E-10** | 1.38E-01 | 3.60E-03 | **1.63E-02** |
| WHR | 0.2 | 2.86E-12 | 1.43E-02 | **2.86E-11** | 0.5 | 7.69E-14 | 2.18E-02 | **7.69E-13** | 1.76E-01 | 2.80E-05 | **6.16E-04** |
| WHR-adjBMI | 0.03 | 2.96E-07 | 1.51E-02 | **2.96E-06** | 0.5 | 1.83E-04 | 1.10E-02 | **1.11E-03** | 1.09E-01 | 7.80E-03 | **2.86E-02** |
| DBP | 0.01 | 9.56E-04 | 3.12E-03 | **9.56E-03** | 0.001 | 4.20E-05 | 1.78E+00 | **4.20E-04** | 1.07E-01 | 3.18E-01 | 4.64E-01 |
| SBP | 0.001 | 2.67E-01 | 9.52E-04 | 9.76E-01 | 0.4 | 9.29E-03 | 3.90E-01 | **3.88E-02** | 1.48E-01 | 2.19E-01 | 4.01E-01 |

Please refer to the legends of Table 1.

Finally, for neuroticism (Table 4), higher PRS of neuroticism was significantly associated with higher fat percentage, FG, HOMA-IR, INS, TC, TG, WHR, DBP and CAD risks. Similar associations were observed when PRS of cardiometabolic traits were used as predictors.

**Table 4.** Polygenic risk score (PRS) analysis between neuroticism and cardiometabolic traits

|  | PRS: neuroticism as outcome |  |  |  | PRS: neuroticism as predictor |  |  |  | LD score regression |  |  |
| --- | --- | --- | --- | --- | --- | --- | --- | --- | --- | --- | --- |
|  | best_p | pval | coef | qval | best_p | pval | coef | qval | rg | pval | qval |
| Adiponectin | 0.05 | 4.29E-01 | -7.90E-04 | 9.09E-01 | 0.3 | 5.29E-03 | -9.21E-03 | **2.41E-02** | -1.00E-01 | 8.30E-02 | 2.80E-01 |
| BMI | 0.001 | 1.64E-02 | -1.63E-02 | 1.64E-01 | 0.001 | 1.39E-01 | -1.18E-02 | 6.71E-01 | -3.49E-02 | 3.57E-01 | 5.61E-01 |
| CAD | 0.01 | 1.45E-08 | 7.40E-03 | **1.35E-07** | 0.3 | 9.37E-10 | 3.16E-02 | **9.37E-09** | 7.83E-02 | 2.77E-02 | 2.03E-01 |
| DM | 0.001 | 1.83E-01 | -1.27E-03 | 8.98E-01 | 0.03 | 4.92E-02 | 2.87E-02 | 2.28E-01 | -4.29E-02 | 4.88E-01 | 6.70E-01 |
| Fat percent | 0.1 | 9.84E-03 | 1.77E-03 | *9.84E-02* | 0.5 | 9.14E-03 | 1.01E-02 | **3.66E-02** | 5.60E-02 | 2.90E-01 | 4.91E-01 |
| FG | 0.03 | 7.86E-02 | 2.92E-03 | 4.71E-01 | 0.1 | 2.45E-03 | 7.94E-03 | **1.18E-02** | 6.10E-02 | 1.73E-01 | 3.80E-01 |
| FG-adjBMI | 0.05 | 5.86E-02 | 2.87E-03 | 3.84E-01 | 0.1 | 2.91E-03 | 7.60E-03 | **9.92E-03** | 6.70E-03 | 8.91E-01 | 8.91E-01 |
| HDL | 0.001 | 3.70E-01 | 4.31E-03 | 9.36E-01 | 0.4 | 2.05E-01 | -4.27E-03 | 8.27E-01 | 7.90E-03 | 8.29E-01 | 8.69E-01 |
| HOMA-IR | 0.03 | 8.99E-05 | 1.06E-02 | **8.99E-04** | 0.5 | 7.92E-04 | 9.71E-03 | **3.51E-03** | 1.13E-01 | 2.41E-01 | 4.81E-01 |
| INS | 0.1 | 4.64E-03 | 3.87E-03 | **4.64E-02** | 0.1 | 3.75E-05 | 9.13E-03 | **3.75E-04** | 1.48E-01 | 1.01E-01 | 2.80E-01 |
| INS-adjBMI | 0.2 | 2.64E-01 | 1.19E-03 | 7.04E-01 | 0.2 | 1.17E-04 | 9.22E-03 | **4.58E-04** | 1.05E-01 | 1.02E-01 | 2.80E-01 |
| LDL | 0.03 | 2.89E-02 | 5.19E-03 | 1.04E-01 | 0.4 | 4.33E-02 | 7.42E-03 | 1.17E-01 | 1.82E-02 | 6.85E-01 | 8.69E-01 |
| Leptin-adjBMI | 0.05 | 3.44E-01 | -1.92E-03 | 9.47E-01 | 0.03 | 3.41E-02 | 1.05E-02 | 1.95E-01 | 1.91E-02 | 8.03E-01 | 8.69E-01 |
| Leptin | 0.01 | 3.05E-01 | 2.49E-03 | 7.73E-01 | 0.4 | 1.46E-01 | 6.67E-03 | 4.07E-01 | 2.12E-02 | 7.56E-01 | 8.69E-01 |
| TC | 0.4 | 3.52E-04 | 5.60E-03 | **1.78E-03** | 0.005 | 1.30E-04 | 2.78E-02 | **1.30E-03** | 4.46E-02 | 2.76E-01 | 4.91E-01 |
| TG | 0.05 | 4.85E-07 | 1.19E-02 | **3.59E-06** | 0.001 | 2.13E-07 | 5.09E-02 | **1.30E-06** | 8.68E-02 | 9.46E-02 | 2.80E-01 |
| WHR | 0.2 | 6.74E-05 | 7.97E-03 | **6.10E-04** | 0.5 | 2.23E-06 | 1.40E-02 | **1.14E-05** | 8.32E-02 | 1.01E-02 | 1.11E-01 |
| WHR-adjBMI | 0.5 | 3.75E-07 | 9.21E-03 | **2.60E-06** | 0.05 | 2.74E-07 | 1.96E-02 | **1.29E-06** | 1.19E-01 | 4.00E-04 | **8.80E-03** |
| DBP | 0.05 | 8.24E-04 | 2.35E-03 | **8.24E-03** | 0.3 | 1.65E-04 | 3.47E-01 | **1.65E-03** | 1.68E-01 | 7.47E-02 | 2.80E-01 |
| SBP | 0.4 | 1.65E-01 | 1.74E-04 | 5.08E-01 | 0.5 | 5.29E-02 | 2.75E-01 | 2.16E-01 | 7.29E-02 | 4.34E-01 | 6.37E-01 |

Please refer to the legends of Table 1.

### LD score regression results

LD score regression in general produced a smaller number of significant results than the polygenic score approach (full results given in Supplementary Table 1). However, most significant results in LDSR were corroborated by significant results from PRS analyses and most significant associations agreed in their directions of effects. For MDD-PGC, nominally significant (*p* < 0.05) associations were observed for TG and WHR (with or without adjustment for BMI) with positive genetic correlations (Table 1). For MDD-CONVERGE, there was nominally significant *negative* genetic correlations with BMI, DM, fat percentage, FG, leptin and WHR. These results are broadly consistent with analyses of polygenic scores, although a few significant results from PRS analyses (e.g. CAD) was not observed in LDSR (Table 2). For depressive symptoms, positive genetic correlations were observed with BMI, CAD, fat percentage, BMI-adjusted leptin, TG and WHR (including BMI-adjusted WHR) and negative genetic correlations with HDL (Table 3). As for neuroticism, nominally significant genetic correlations were observed for CAD and WHR (including BMI-adjusted WHR) (Table 4).

Of note, the intercept from LDSR for MDD-PGC, depressive symptoms and neuroticism were all non-significant (*p* > 0.05), except for one pair of traits (TG with MDD-PGC; *p* = 0.049), which is within expectations (we expect 54*0.05 = 2.7 significant associations by chance) (Supplementary Table 1). The results suggest no or only minor overlap between the samples. LD score regression was not performed for MDD-CONVERGE as the sample is completely separate from other studies.

### Identification of shared genetic variants and Pathway/GO term over-representation analyses

The top genes shared by MDD and cardiometabolic traits are shown in Tables 5. To facilitate interpretation, we have performed LD-clumping (with an R-squared threshold of 0.1 and a window-size of 250 kb) on the shared SNPs using PLINK^54^ and mapped the SNPs to genes using BioMart. As discussed above, for SNPs mapping to more than one gene, we only show the gene associated with the worst variant consequence. For easy visualization of results, we also prepared a crude summary table which includes the number of times each gene appears on our top list (defined as tdr_11_> 0.5) across the 20 cardiometabolic traits and the corresponding average rank. Similar summary tables were also created for pathway and GO terms analyses. Supplementary Tables 2-5 show all LD-clumped SNPs with tdr_11_> 0.5 that map to at least one gene.

**Table 5.**
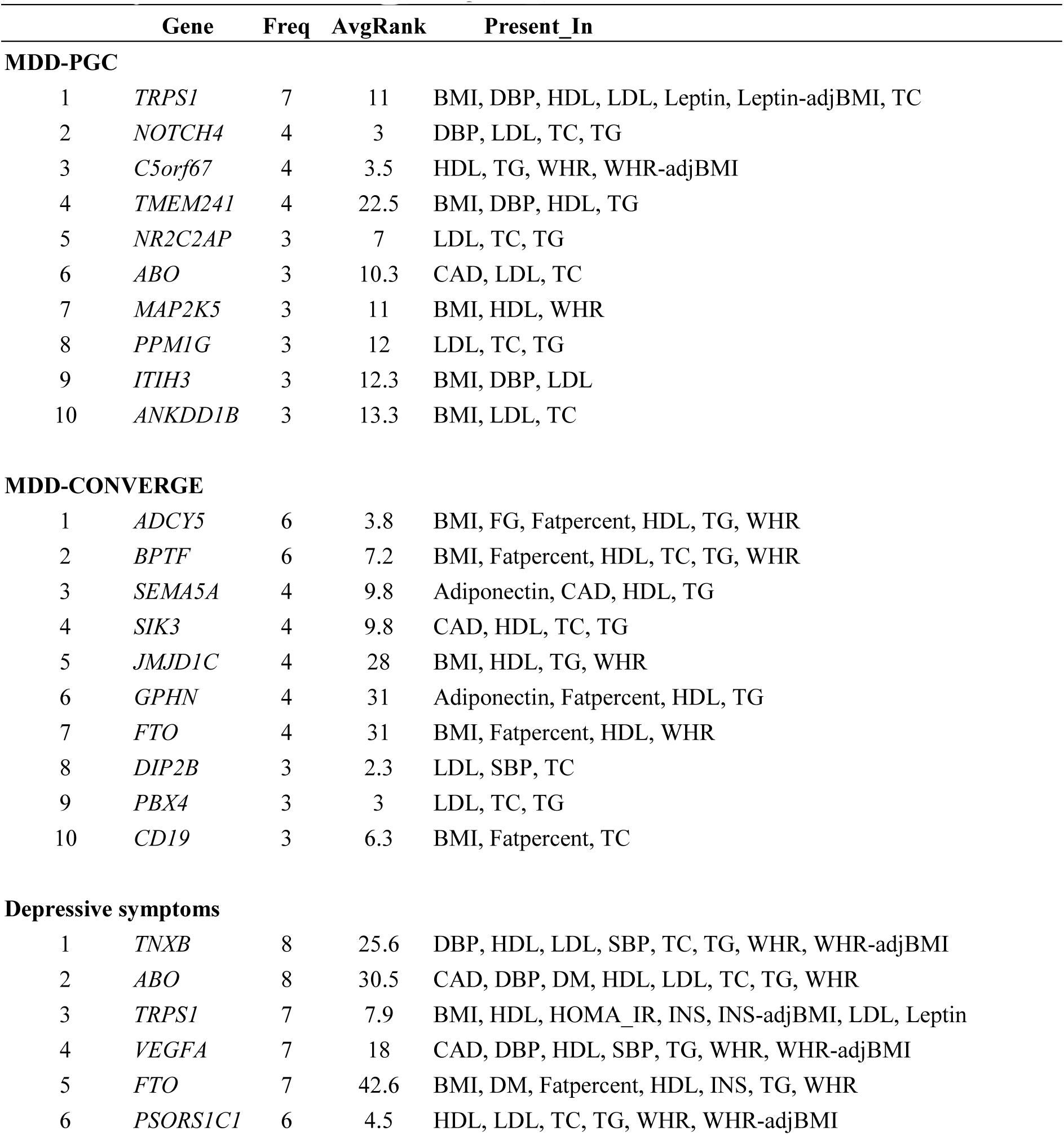
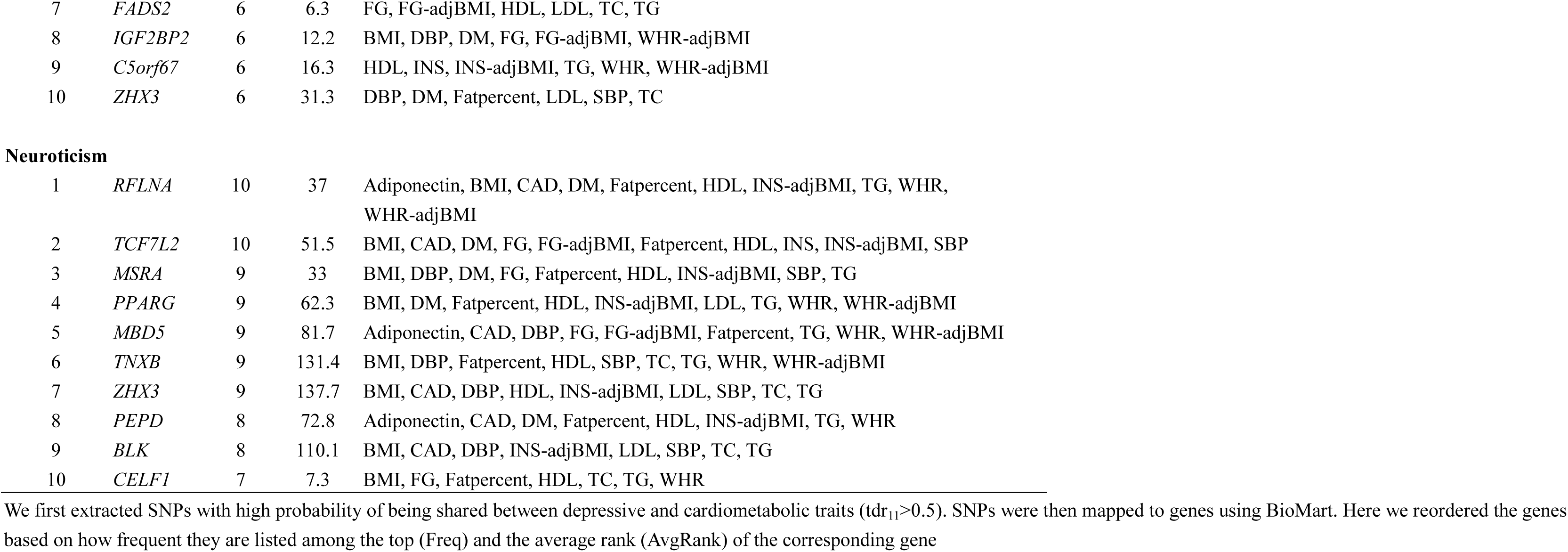
Top genes shared between depression phenotypes and cardiometabolic traits

|  | Gene | Freq | AvgRank | Present_In |
| --- | --- | --- | --- | --- |
| **MDD-PGC** |  |  |  |  |
| 1 | *TRPS1* | 7 | 11 | BMI, DBP, HDL, LDL, Leptin, Leptin-adjBMI, TC |
| 2 | *NOTCH4* | 4 | 3 | DBP, LDL, TC, TG |
| 3 | *C5orf67* | 4 | 3.5 | HDL, TG, WHR, WHR-adjBMI |
| 4 | *TMEM241* | 4 | 22.5 | BMI, DBP, HDL, TG |
| 5 | *NR2C2AP* | 3 | 7 | LDL, TC, TG |
| 6 | *ABO* | 3 | 10.3 | CAD, LDL, TC |
| 7 | *MAP2K5* | 3 | 11 | BMI, HDL, WHR |
| 8 | *PPM1G* | 3 | 12 | LDL, TC, TG |
| 9 | *ITIH3* | 3 | 12.3 | BMI, DBP, LDL |
| 10 | *ANKDD1B* | 3 | 13.3 | BMI, LDL, TC |
| **MDD-CONVERGE** |  |  |  |  |
| 1 | *ADCY5* | 6 | 3.8 | BMI, FG, Fatpercent, HDL, TG, WHR |
| 2 | *BPTF* | 6 | 7.2 | BMI, Fatpercent, HDL, TC, TG, WHR |
| 3 | *SEMA5A* | 4 | 9.8 | Adiponectin, CAD, HDL, TG |
| 4 | *SIK3* | 4 | 9.8 | CAD, HDL, TC, TG |
| 5 | *JMJD1C* | 4 | 28 | BMI, HDL, TG, WHR |
| 6 | *GPHN* | 4 | 31 | Adiponectin, Fatpercent, HDL, TG |
| 7 | *FTO* | 4 | 31 | BMI, Fatpercent, HDL, WHR |
| 8 | *DIP2B* | 3 | 2.3 | LDL, SBP, TC |
| 9 | *PBX4* | 3 | 3 | LDL, TC, TG |
| 10 | *CD19* | 3 | 6.3 | BMI, Fatpercent, TC |
| **Depressive symptoms** |  |  |  |  |
| 1 | *TNXB* | 8 | 25.6 | DBP, HDL, LDL, SBP, TC, TG, WHR, WHR-adjBMI |
| 2 | *ABO* | 8 | 30.5 | CAD, DBP, DM, HDL, LDL, TC, TG, WHR |
| 3 | *TRPS1* | 7 | 7.9 | BMI, HDL, HOMA_IR, INS, INS-adjBMI, LDL, Leptin |
| 4 | *VEGFA* | 7 | 18 | CAD, DBP, HDL, SBP, TG, WHR, WHR-adjBMI |
| 5 | *FTO* | 7 | 42.6 | BMI, DM, Fatpercent, HDL, INS, TG, WHR |
| 6 | *PSORS1C1* | 6 | 4.5 | HDL, LDL, TC, TG, WHR, WHR-adjBMI |
| 7 | *FADS2* | 6 | 6.3 | FG, FG-adjBMI, HDL, LDL, TC, TG |
| 8 | *IGF2BP2* | 6 | 12.2 | BMI, DBP, DM, FG, FG-adjBMI, WHR-adjBMI |
| 9 | *C5orf67* | 6 | 16.3 | HDL, INS, INS-adjBMI, TG, WHR, WHR-adjBMI |
| 10 | *ZHX3* | 6 | 31.3 | DBP, DM, Fatpercent, LDL, SBP, TC |
| Neuroticism |  |  |  |  |
| 1 | *RFLNA* | 10 | 37 | Adiponectin, BMI, CAD, DM, Fatpercent, HDL, INS-adjBMI, TG, WHR, WHR-adjBMI |
| 2 | *TCF7L2* | 10 | 51.5 | BMI, CAD, DM, FG, FG-adjBMI, Fatpercent, HDL, INS, INS-adjBMI, SBP |
| 3 | *MSRA* | 9 | 33 | BMI, DBP, DM, FG, Fatpercent, HDL, INS-adjBMI, SBP, TG |
| 4 | *PPARG* | 9 | 62.3 | BMI, DM, Fatpercent, HDL, INS-adjBMI, LDL, TG, WHR, WHR-adjBMI |
| 5 | *MBD5* | 9 | 81.7 | Adiponectin, CAD, DBP, FG, FG-adjBMI, Fatpercent, TG, WHR, WHR-adjBMI |
| 6 | *TNXB* | 9 | 131.4 | BMI, DBP, Fatpercent, HDL, SBP, TC, TG, WHR, WHR-adjBMI |
| 7 | *ZHX3* | 9 | 137.7 | BMI, CAD, DBP, HDL, INS-adjBMI, LDL, SBP, TC, TG |
| 8 | *PEPD* | 8 | 72.8 | Adiponectin, CAD, DM, Fatpercent, HDL, INS-adjBMI, TG, WHR |
| 9 | *BLK* | 8 | 110.1 | BMI, CAD, DBP, INS-adjBMI, LDL, SBP, TC, TG |
| 10 | *CELF1* | 7 | 7.3 | BMI, FG, Fatpercent, HDL, TC, TG, WHR |

We first extracted SNPs with high probability of being shared between depressive and cardiometabolic traits ( $\text{tdr}_{11} > 0.5$ ). SNPs were then mapped to genes using BioMart. Here we reordered the genes based on how frequent they are listed among the top (Freq) and the average rank (AvgRank) of the corresponding gene

To explore whether the genes shared by depression and cardiometabolic traits share specific pathways and functional features, we performed pathway and GO enrichment analysis using ConsensusPathDB. Tables 6 and 7 summarize the pathway enrichment results for the four depression traits under study. Here we mainly focused on the pathways derived from SNPs that increase (or decrease) the risks of depression and CV abnormalities at the same time (note that for HDL and adiponectin, lower levels are associated with higher CV risks). For MDD-CONVERGE, since we observed unexpected inverse polygenic associations with many CV risk factors, we presented the top enriched pathways derived from SNPs having opposite effects on depression and CV risks. The full results of pathway and GO term enrichment, together with summary tables of the most frequently enriched pathways across different traits, are available in Supplementary Tables 6-13.

**Table 6.**
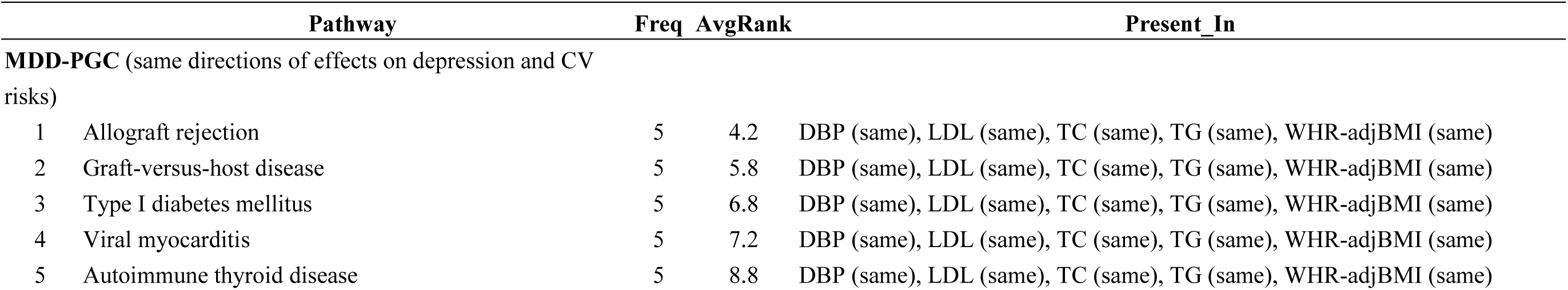
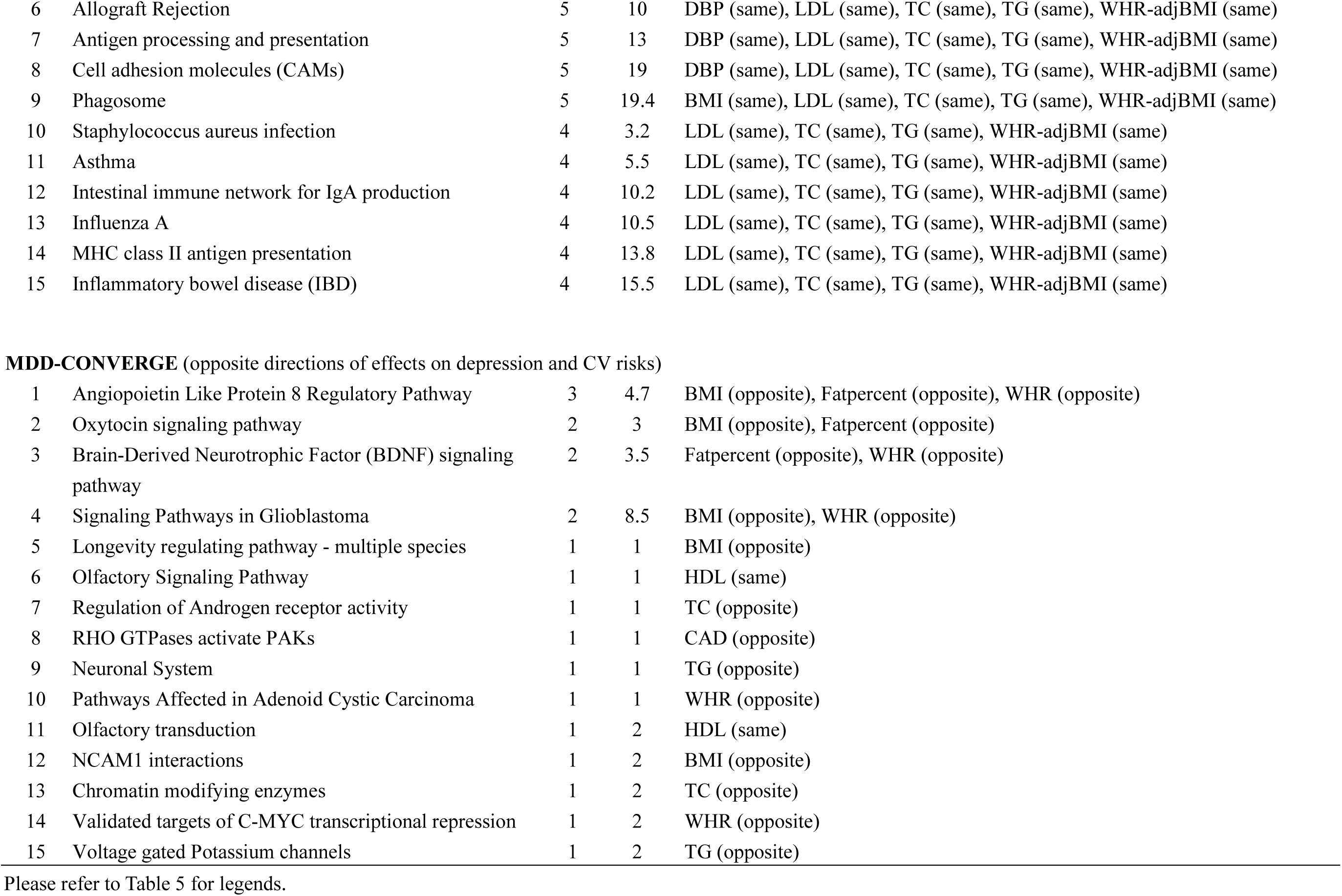
Top pathways enriched among genes shared by MDD-PGC and MDD-CONVEGE with cardiometabolic traits

|  | Pathway | Freq | AvgRank | Present_In |
| --- | --- | --- | --- | --- |
| **MDD-PGC** (same directions of effects on depression and CV risks) |  |  |  |  |
| 1 | Allograft rejection | 5 | 4.2 | DBP (same), LDL (same), TC (same), TG (same), WHR-adjBMI (same) |
| 2 | Graft-versus-host disease | 5 | 5.8 | DBP (same), LDL (same), TC (same), TG (same), WHR-adjBMI (same) |
| 3 | Type I diabetes mellitus | 5 | 6.8 | DBP (same), LDL (same), TC (same), TG (same), WHR-adjBMI (same) |
| 4 | Viral myocarditis | 5 | 7.2 | DBP (same), LDL (same), TC (same), TG (same), WHR-adjBMI (same) |
| 5 | Autoimmune thyroid disease | 5 | 8.8 | DBP (same), LDL (same), TC (same), TG (same), WHR-adjBMI (same) |
| 6 | Allograft Rejection | 5 | 10 | DBP (same), LDL (same), TC (same), TG (same), WHR-adjBMI (same) |
| 7 | Antigen processing and presentation | 5 | 13 | DBP (same), LDL (same), TC (same), TG (same), WHR-adjBMI (same) |
| 8 | Cell adhesion molecules (CAMs) | 5 | 19 | DBP (same), LDL (same), TC (same), TG (same), WHR-adjBMI (same) |
| 9 | Phagosome | 5 | 19.4 | BMI (same), LDL (same), TC (same), TG (same), WHR-adjBMI (same) |
| 10 | Staphylococcus aureus infection | 4 | 3.2 | LDL (same), TC (same), TG (same), WHR-adjBMI (same) |
| 11 | Asthma | 4 | 5.5 | LDL (same), TC (same), TG (same), WHR-adjBMI (same) |
| 12 | Intestinal immune network for IgA production | 4 | 10.2 | LDL (same), TC (same), TG (same), WHR-adjBMI (same) |
| 13 | Influenza A | 4 | 10.5 | LDL (same), TC (same), TG (same), WHR-adjBMI (same) |
| 14 | MHC class II antigen presentation | 4 | 13.8 | LDL (same), TC (same), TG (same), WHR-adjBMI (same) |
| 15 | Inflammatory bowel disease (IBD) | 4 | 15.5 | LDL (same), TC (same), TG (same), WHR-adjBMI (same) |
| **MDD-CONVERGE** (opposite directions of effects on depression and CV risks) |  |  |  |  |
| 1 | Angiopoietin Like Protein 8 Regulatory Pathway | 3 | 4.7 | BMI (opposite), Fatpercent (opposite), WHR (opposite) |
| 2 | Oxytocin signaling pathway | 2 | 3 | BMI (opposite), Fatpercent (opposite) |
| 3 | Brain-Derived Neurotrophic Factor (BDNF) signaling pathway | 2 | 3.5 | Fatpercent (opposite), WHR (opposite) |
| 4 | Signaling Pathways in Glioblastoma | 2 | 8.5 | BMI (opposite), WHR (opposite) |
| 5 | Longevity regulating pathway - multiple species | 1 | 1 | BMI (opposite) |
| 6 | Olfactory Signaling Pathway | 1 | 1 | HDL (same) |
| 7 | Regulation of Androgen receptor activity | 1 | 1 | TC (opposite) |
| 8 | RHO GTPases activate PAKs | 1 | 1 | CAD (opposite) |
| 9 | Neuronal System | 1 | 1 | TG (opposite) |
| 10 | Pathways Affected in Adenoid Cystic Carcinoma | 1 | 1 | WHR (opposite) |
| 11 | Olfactory transduction | 1 | 2 | HDL (same) |
| 12 | NCAM1 interactions | 1 | 2 | BMI (opposite) |
| 13 | Chromatin modifying enzymes | 1 | 2 | TC (opposite) |
| 14 | Validated targets of C-MYC transcriptional repression | 1 | 2 | WHR (opposite) |
| 15 | Voltage gated Potassium channels | 1 | 2 | TG (opposite) |

Please refer to Table 5 for legends.

**Table 7.**
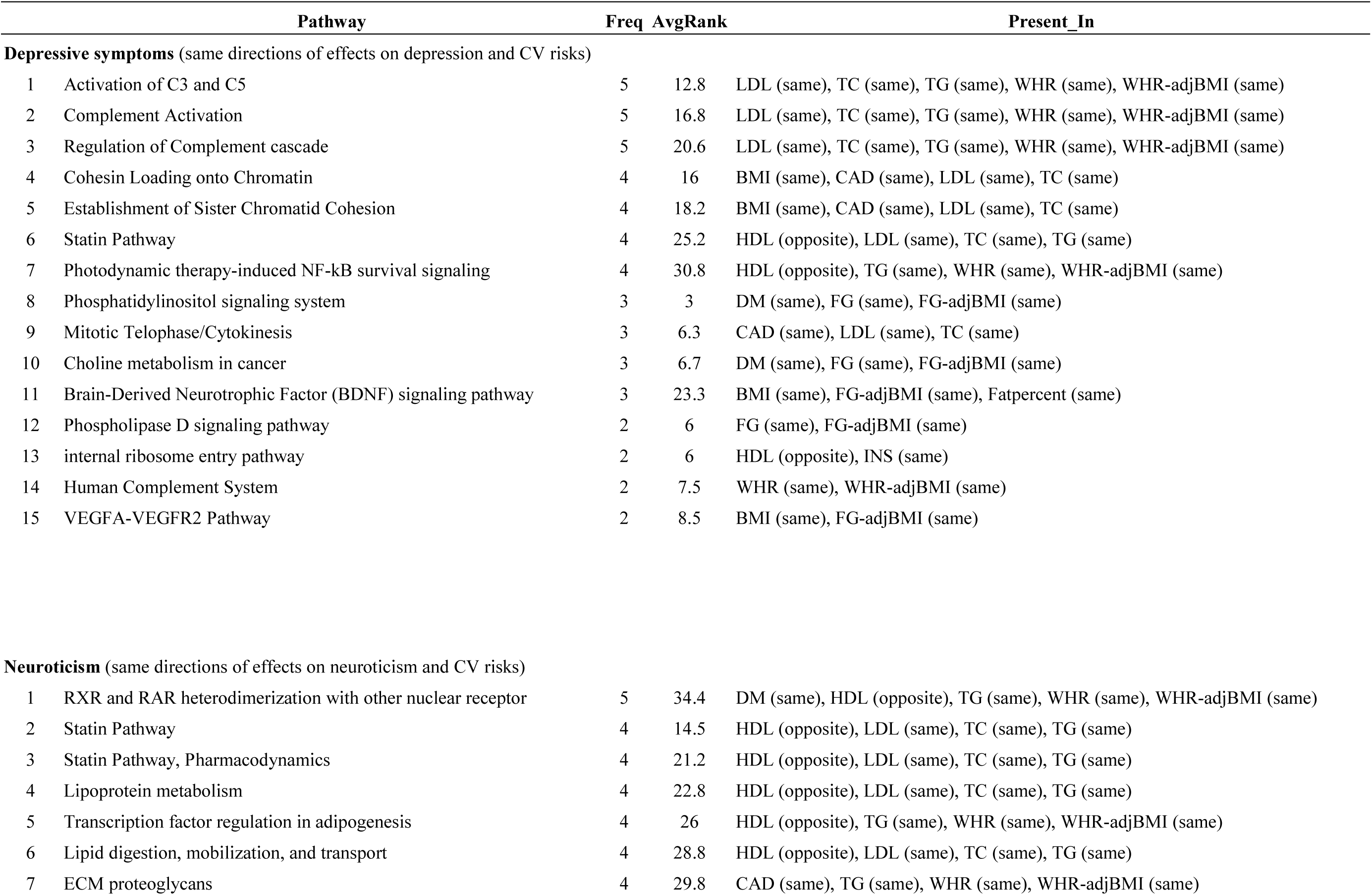
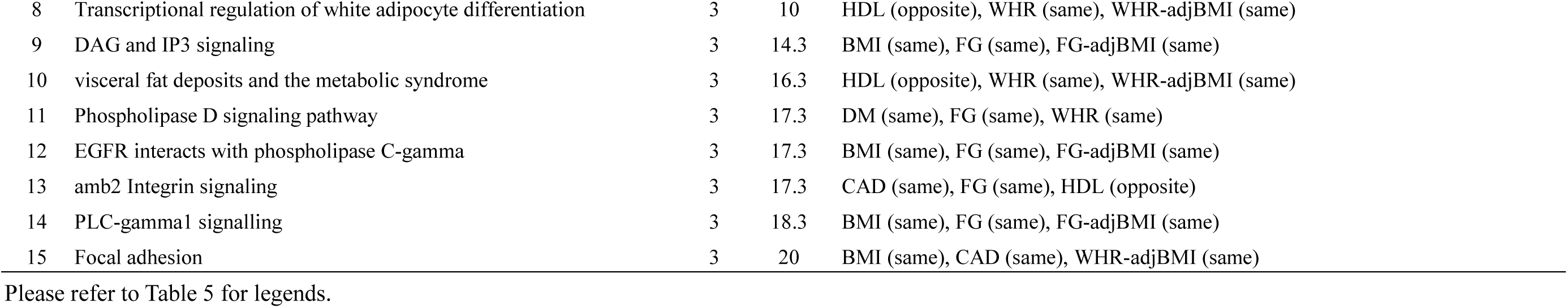
Top pathways enriched among genes shared by depressive symptoms and neuroticism with cardiometabolic traits

For MDD-PGC, we observed that immune- or inflammation-related pathways were frequently enriched across numerous cardiometabolic traits; genes within the HLA region contributed to many of these pathways. Some other interesting pathways enriched included biological oxidation (enriched in CAD, SBP and DBP), chaperonin-mediated protein folding (BMI, LDL, TC) and purine metabolism (SBP, DBP) (Table 6 and Supplementary Table 7).

For MDD-CONVERGE, when we considered pathways derived from SNPs that were associated with increases in both depression and CV risks, several pathways were often enriched such as metabolism of vitamins and cofactors, GABAergic synapse and activation of gene expression by SREBF (Supplementary Table 9). On the other hand, when the focus was on SNPs conferring *opposite* risks for depression and CV diseases, the Angiopoietin Like Protein 8 (ANGPTL8) regulatory pathway, oxytocin signaling pathway and the Brain-Derived Neurotrophic Factor (BDNF) pathway were listed among the top (Table 6 and Supplementary Table 9).

Concerning depressive symptoms, some of the prioritized pathways included complement activation pathways (LDL, TC, TG, WHR; contributed by the genes *C2*, *C7* and *CFB*), the statin pathway (LDL, TC, TG, HDL), phosphatidylinositol signaling (DM, FG), brain-derived neurotrophic factor (BDNF) signaling (BMI, FG, fat percentage) and the VEGF pathways (BMI, FG), among others (Table 7 and Supplementary Table 11).

For neuroticism, some of the most often enriched pathways included ‘RXR and RAR heterodimerization with other nuclear receptor’ (DM, TG, WHR, HDL), the statin pathway and other lipid metabolism or adipogenesis pathways (LDL, TC, TG, HDL, CAD, DM), ECM proteoglycans (CAD, TG, WHR), DAG and IP3 signaling (BMI, FG), phospholipase D signaling (DM, FG, WHR) and EGFR signaling (BMI, FG) (Table 7 and Supplementary Table 13).

### Drug enrichment analysis

Here our focus is on finding drugs with repositioning potential for comorbid depression and CV diseases; hence the drug enrichment analysis was performed with genetic variants that exert the *same* directions of effects on depression and CV risks simultaneously. The drug enrichment results are shown in Table 8 and Supplementary Tables 14-17. Some of the interesting hits included the anti-oxidant glutathione, the xanthine oxidase inhibitor allopurinol and statins such as rosuvastatin and simvastatin. For MDD-CONVERGE, we still focused on analysis of SNPs that raise depression and CV risks at the same time. Some of the interesting top-listed candidates included several statins, glycine, bupropion and verapamil. For SSGAC-DS, some commonly enriched drugs included lipase, several statins, bupropion, fenofibrate, glutathione, among others. For neuroticism, some top hits included insulin, glutathione, bupropion, statins, fluphenazine and gemfibrozil.

**Table 8.**
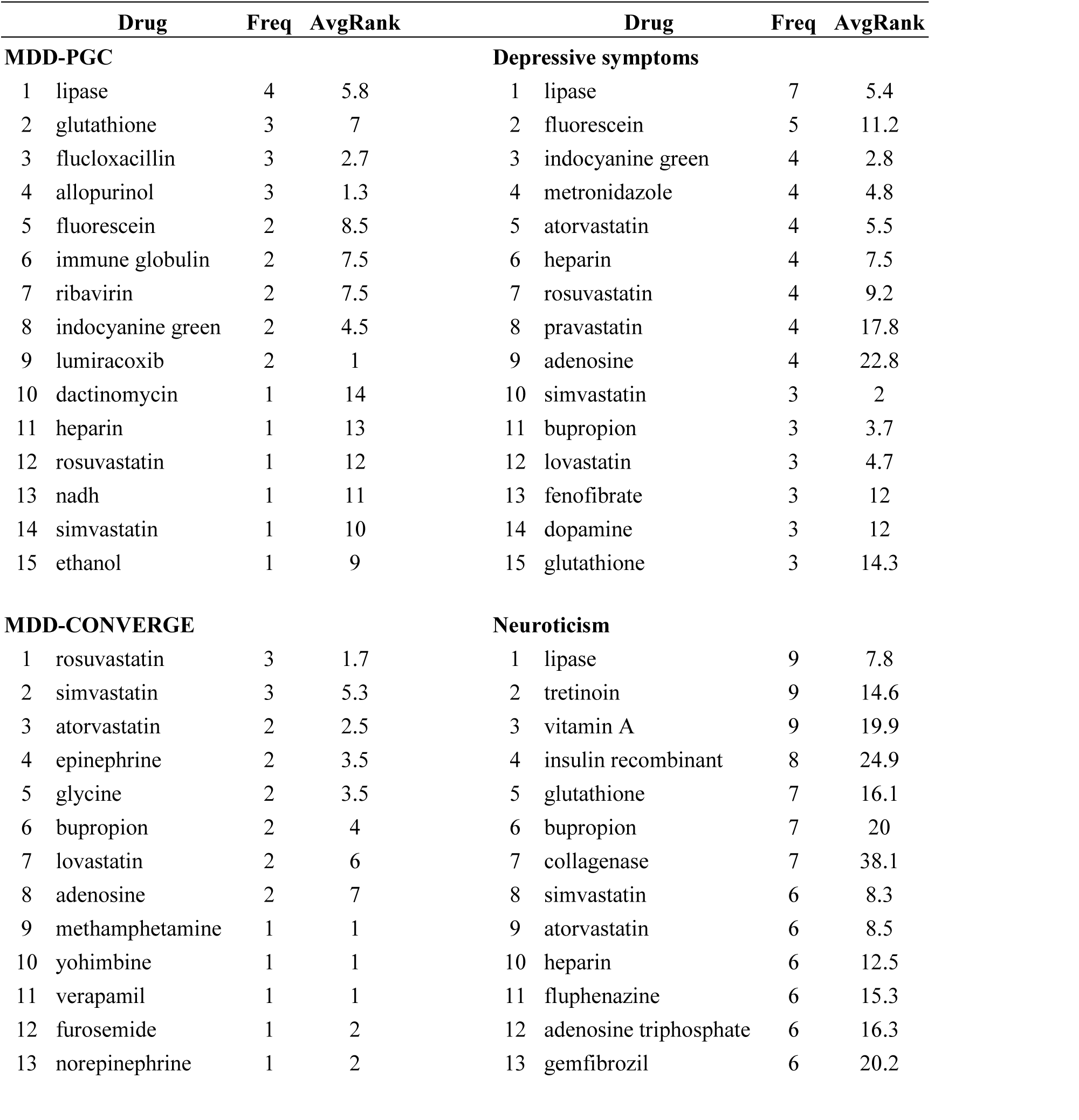
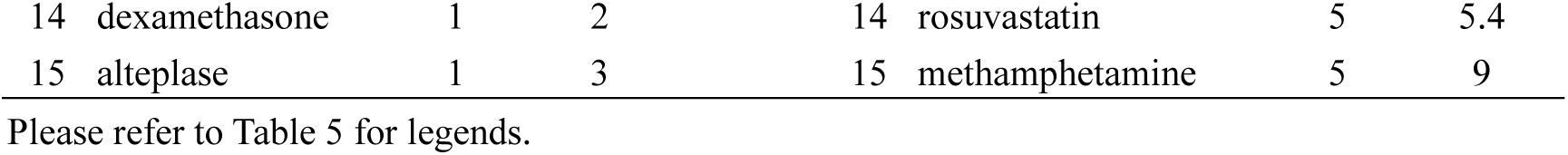
Drugs enriched among the top shared genes between depression and cardiometabolic traits

| Drug | Freq | AvgRank | Drug | Freq | AvgRank |
| --- | --- | --- | --- | --- | --- |
| **MDD-PGC** |  |  | **Depressive symptoms** |  |  |
| 1 lipase | 4 | 5.8 | 1 lipase | 7 | 5.4 |
| 2 glutathione | 3 | 7 | 2 fluorescein | 5 | 11.2 |
| 3 flucloxacillin | 3 | 2.7 | 3 indocyanine green | 4 | 2.8 |
| 4 allopurinol | 3 | 1.3 | 4 metronidazole | 4 | 4.8 |
| 5 fluorescein | 2 | 8.5 | 5 atorvastatin | 4 | 5.5 |
| 6 immune globulin | 2 | 7.5 | 6 heparin | 4 | 7.5 |
| 7 ribavirin | 2 | 7.5 | 7 rosuvastatin | 4 | 9.2 |
| 8 indocyanine green | 2 | 4.5 | 8 pravastatin | 4 | 17.8 |
| 9 lumiracoxib | 2 | 1 | 9 adenosine | 4 | 22.8 |
| 10 dactinomycin | 1 | 14 | 10 simvastatin | 3 | 2 |
| 11 heparin | 1 | 13 | 11 bupropion | 3 | 3.7 |
| 12 rosuvastatin | 1 | 12 | 12 lovastatin | 3 | 4.7 |
| 13 nadh | 1 | 11 | 13 fenofibrate | 3 | 12 |
| 14 simvastatin | 1 | 10 | 14 dopamine | 3 | 12 |
| 15 ethanol | 1 | 9 | 15 glutathione | 3 | 14.3 |
| **MDD-CONVERGE** |  |  | **Neuroticism** |  |  |
| 1 rosuvastatin | 3 | 1.7 | 1 lipase | 9 | 7.8 |
| 2 simvastatin | 3 | 5.3 | 2 tretinoin | 9 | 14.6 |
| 3 atorvastatin | 2 | 2.5 | 3 vitamin A | 9 | 19.9 |
| 4 epinephrine | 2 | 3.5 | 4 insulin recombinant | 8 | 24.9 |
| 5 glycine | 2 | 3.5 | 5 glutathione | 7 | 16.1 |
| 6 bupropion | 2 | 4 | 6 bupropion | 7 | 20 |
| 7 lovastatin | 2 | 6 | 7 collagenase | 7 | 38.1 |
| 8 adenosine | 2 | 7 | 8 simvastatin | 6 | 8.3 |
| 9 methamphetamine | 1 | 1 | 9 atorvastatin | 6 | 8.5 |
| 10 yohimbine | 1 | 1 | 10 heparin | 6 | 12.5 |
| 11 verapamil | 1 | 1 | 11 fluphenazine | 6 | 15.3 |
| 12 furosemide | 1 | 2 | 12 adenosine triphosphate | 6 | 16.3 |
| 13 norepinephrine | 1 | 2 | 13 gemfibrozil | 6 | 20.2 |

|  |  |  |  |  |  |  |  |
| --- | --- | --- | --- | --- | --- | --- | --- |
| 14 | dexamethasone | 1 | 2 | 14 | rosuvastatin | 5 | 5.4 |
| 15 | alteplase | 1 | 3 | 15 | methamphetamine | 5 | 9 |

Please refer to Table 5 for legends.

## Discussion

In this study, we discovered significant polygenic sharing between depression-related phenotypes with multiple cardiometabolic traits, and inferred pathways that were enriched among the shared genes. We also explored drugs with the potential of being repositioned for the comorbid disorders. The results of each part of the analyses are discussed below.

### Polygenic associations of various depression-related phenotypes with cardiometabolic traits

Overall we have discovered significant polygenic associations of 3 depression or depression-related phenotypes (MDD-PGC, depressive symptoms, neuroticism) with increased cardiometabolic risks spanning multiple traits; however, for MDD-CONVERGE, we observed an opposite trend of reduced cardiometabolic risks. To our knowledge, this is the first comprehensive survey of polygenic overlap and shared genetic variants between depression traits and a whole spectrum of cardiometabolic abnormalities, including lipid- and glucose-related traits, blood pressure, anthropometric and adiposity measures, CAD, DM, as well as adipokine abnormalities.

#### Neuroticism and cardiometabolic abnormalities

As introduced earlier, numerous studies have reported raised CV risks in patients with depression or higher depressive symptoms, as reviewed elsewhere^5,55^. Nevertheless, far less studies have investigated possible shared genetic bases of these disorders. A recent study by Gale et al.^56^ investigated genetic sharing between neuroticism and several physical and mental health traits. Polygenic score analysis was performed on 108038 UK biobank (UKB) participants while LDSR was conducted separately on both UKB and the Genetics of Personality Consortium (GPC). The SSGAC summary statistics we used in this study were derived from meta-analysis of both datasets. Therefore, there was a partial overlap of samples with the analysis by Gale et al.^56^, but the current work encompassed a larger variety of CV traits (20 vs 5). Similar to Gale et al.^56^, we found a significant polygenic association with CAD and but no association with DM. An inverse relationship of neuroticism was found with BMI in the previous study with the UKB sample alone^56^, but we did not observe any associations using combined results from UKB and GPC. It is possible that the differences in survey instruments and heterogeneity of study samples may contribute to this difference.

Several epidemiological studies have studied the relationship between neuroticism and cardiovascular diseases, but the results have been variable (*e.g.* ref.^57-63^). The current study revealed positive polygenic associations with lipid traits as well as higher fat percentage and WHR, although there was no significant association with BMI. Interestingly, we observed positive polygenic relationships with insulin resistance and hyperglycemia, but no significant connection with DM. It remains to be investigated whether this is the case in practice, and whether individuals having more extreme scores on neuroticism will harbor higher diabetic risks. Nonetheless, the genetic bases of glucose levels and DM might not completely overlap. For example, Merino et al. reported that the polygenic score of 12 variants having strong associations with FG did not significantly predict the risk of type 2 DM^64^ and some SNPs may have differential effects on quantitative glycemic traits and the risk of frank diabetes^65,66^; a more detailed review was given by Marullo et al.^67^. On the other hand, a highly significant association was observed with CAD, providing strong support for a positive association and shared genetics of the two traits. Neuroticism is a well-established risk factor for major depression and anxiety disorders^68^; it remains to be studied whether the raised CV risks conferred by neuroticism is mediated by these associated conditions or neuroticism is itself an independent risk factor. Interestingly, a recent study demonstrated a synergistic interaction between neuroticism and MDD on the risks of CV diseases^7^.

#### MDD-PGC and SSGAC depressive symptoms study: relationship with cardiometabolic abnormalities

It is worth noting both the MDD-PGC and SSAGC-DS samples did not stratify depression patients according to subtypes or other clinical features. The SSGAC study sample also contained a substantial proportion of UKB participants whose depressive symptoms were measured by self-reported questionnaires^23^. Previous epidemiological studies in this area have also been based on both clinical diagnosis as well as symptom questionnaires^55^. Although higher^69^, lower^16^ or similar^70^ effect sizes have all been reported with questionnaire-based studies, the overall trend was similar, in that depression increases CV risks. Here we observed largely consistent polygenic associations with cardiometabolic traits for both MDD-PGC and the SSGAC-DS. With the exception of lowered adiponectin in MDD-PGC, all significant associations in MDD-PGC were also observed in SSGAC-DS. A wider range of cardiometabolic traits were significantly associated with SSGAC-DS, which may be explained by its larger sample size (∼ 10 times of MDD-PGC). Notably, we observed polygenic associations across almost the entire range of CV risk factors for SSGAC-DS, and all associations were in the direction of increased CV risks. This observation provides strong support for a shared genetic basis and pathophysiology between depression and CV risk factors and diseases.

There were a few unexpected polygenic associations observed for MDD-PGC. Lower BMI-adjusted leptin was observed to be associated with depression, however higher leptin levels, indicating leptin resistance, are associated with CV risks. Unadjusted leptin however did not show any associations. We suspected that this may be due to ‘collider bias’^71^, in which the adjustment of a covariate positively associated with both the predictor and the outcome can sometimes lead to unexpected inverse associations even when such relationships are absent (*e.g.*^72^). Nevertheless, in this study we did not observe any other similar examples indicating this collider bias. In most cases the BMI-adjusted estimates were concordant with the unadjusted estimates in PRS analyses. Although this is a potential source of bias, we still included BMI-adjusted CV traits in our analyses as this information is pottentially important in showing whether the trait is a significant risk factor independent of BMI, and that there are no current no good methods to circumvent this problem^71^. Another slightly unexpected finding is that PRS of FG was inversely associated with depression. Previous studies have reported more severe depression symptoms among type 2 DM patients with self-reported hypoglycemia^73^, however the underlying mechanism is unclear. It should be noted that there is incomplete overlap of the genetic bases of glucose levels and frank DM^67^.

#### MDD-CONVERGE (severe melancholic depression) and cardiometabolic abnormalities

Surprisingly, we observed *inverse* polygenic associations of a number of cardiometabolic abnormalities, including DM and CAD, with MDD-CONVERGE. The MDD-CONVERGE sample was mainly composed of severe depression patients with recurrent disease episodes. Notably, 85% of the cases met the DSM-IV criteria for melancholia. The sample was recruited from China and contained only women, with cases predominantly selected from hospitalized patients^74^. Furthermore, as remarked by the study authors, there is probably a reluctance or lower awareness to report depression or to seek medical attention for MDD in China^21^; this is reflected by a ∼5 fold lower prevalence of MDD in China^75^ than that in the US^76^. Those who were actually diagnosed of MDD and hospitalized may therefore represent a particularly severe group of patients.

The proportion of MDD patients with melancholic features is ∼ 23.5% according a large study by McGrath et al.^77^. Given the sample selection method and possible reluctance to report MDD in China, the overall proportion of MDD patients with comparable severity and clinical features to the CONVERGE sample is probably much less than 23.5% (say below one-fifth or even lower). As this subgroup of patients represent a minority, this may partially explain why meta-analyses of epidemiological studies have consistently found an *increased* risk of CV disorders in MDD patients. Even if a subset of patients have reduced CV risks, they may be overwhelmed by the rest of the sample harboring heightened risks.

While the inverse polygenic associations are intriguing, the current study does *not* immediately signify that severe melancholic depression patients must have lower CV risks *in practice*. This is because the focus of the current study is on shared *genetic* factors of depression and CV diseases, but we did not directly address other behavioral factors that can lead to the comorbidities. For example, depressed patients are less likely to adhere to drug treatments and risk-factor modification interventions^78^ and may be more likely to be physically inactive^79^. In addition, severe depression patients may have high chances of receiving augmentation therapy, for example by atypical antipsychotics^80^ or lithium^81^, which may also contribute to weight gain and cardiometabolic abnormalities^82,83^. However, given similar risk factor profiles, this study suggests that patients with non-melancholic depression might be at higher CV risks than their melancholic counterparts; greater awareness and more intense monitoring of CV comorbidities in non-melancholic patients might be warranted. Another implication is that if shared pathophysiology is unimportant for CV risks in melancholic patients, then behavioral interventions (*e.g.* program to foster treatment compliance) may be sufficient to ameliorate CV risks in these patients, as opposed to non-melancholic depression. This hypothesis requires verification in future studies.

To our knowledge, this is the first study showing *inverse* polygenic associations of melancholic depression with cardiometabolic risks. Relatively few epidemiological studies have addressed CV comorbidities with consideration of depression subtypes. For example, Seppala et al.^84^ found that the prevalence of metabolic syndrome (MetS) is about 2-fold higher within atypical MDD patients as compared to controls; there was no significant association with melancholic MDD. Another study also reported association of atypical depression with increased prevalence of overweight, DM and MetS, while melancholic depression was associated with significantly lower prevalence of overweight^85^. Lamers et al.^18^ revealed that levels of inflammatory markers, BMI, TG, waist circumference were higher in atypical MDD patients than melancholic patients or controls. In fact SBP and BMI were significantly *lower* in melancholic patients compared to controls in the study. Yet a more recent study^20^ reported that only the atypical subtype of MDD was prospectively associated with an elevated incidence of MetS and a steeper increase in FG level. Notably, these associations were found to be independent of eating behaviors, psychiatric comorbidities, lifestyle factors, or baseline adipokine concentrations and inflammatory marker levels^20^. Taken together observations from previous epidemiological studies were broadly consistent with our findings that CV risks differ by depression subtypes, and non-melancholic patients were generally at higher risks.

In summary, our findings support a genetic basis for the heterogeneity of depression comorbidities, and that subtyping depression patients might help to better delineate patients’ CV risks, which may have implications in targeted prevention of CV events for patients.

### Shared genetic variants and pathways

We discovered numerous shared genetic variants across the 20 cardiometabolic traits. Some interesting genes listed included *FTO*, which is a well-known gene in controlling weight and energy balance^86^; *NOTCH4*, a member of the NOTCH signaling pathway which is involved in various developmental and homeostatic processes^87^, with mutations being associated with schizophrenia^88^; *TCF7L2,* a transcription factor of key role of Wnt signaling pathway that is strongly associated with diabetes^89^; *PPARG*, a target for anti-diabetic agents (notably the PPAR-gamma agonist pioglitazone has been shown to be efficacious in depression^90^); *VEGFA*, a member of the vascular endothelial growth factor family which acts on endothelial cells with myriad functions^91^ but is also implicated neuropsychiatric disorders^92^.

Due to the wide range of traits studied and the large number of shared variants, we shall not discuss each gene in detail. Here we briefly highlight several interesting pathways that are shared between depression and cardiometabolic traits inferred from shared SNPs. Notably, numerous immune- or inflammation-related pathways were listed among the top for MDD-PGC and SSGAC-DS. Increased inflammation has long been postulated as one of the mechanisms linking the two kinds of disorders^93^. For example, inflammatory markers such as tumor necrosis factor, interleukin-6 (IL-6) and C-reactive protein (CRP) were elevated in MDD as shown in meta-analyses^94,95^. Elevation of some of these markers have also found in comorbid depression and metabolic syndrome^96^ as well as CAD^97^. It has also been suggested that adiposity may mediate the increased inflammatory response seen in depression^98^. There is some evidence that antidepressants reduce inflammatory responses^99^. However, increased inflammation may not be observed in every depressive patient^55^, and further studies are required to elucidate the subgroup of patients most vulnerable to the effects of increased inflammatory response.

Interestingly, the statin pathway was highly ranked for depressive symptoms and neuroticism. Consistent with our results, statins have been found to exert many pleiotropic beneficial effects on cardiovascular diseases, including anti-inflammatory actions, beyond their LDL cholesterol-lowering effects ^100^. Recently, some researchers started to study its therapeutic potential on depression. A Swedish national cohort study reported that statin possibly reduced risk of depression in individuals over the age of 40 ^103^. It was suggested that concomitant treatment with selective serotonin reuptake inhibitor (SSRI) and statin might have superior antidepressant effect than SSRI treatment alone ^104^. A Korean group also successfully showed an antidepressant action of statin in patients with acute coronary syndrome (ACS) ^105^. Nevertheless, studies have also reported increased depressive symptoms and risk of suicide among statin users as reviewed by You et al.^106^. Further studies are required to clarify which kinds of factors (*e.g.* type of statins used, duration of treatment, concomitant medications, patient characteristics) will influence the action of statins on depressive symptoms.

Some other noteworthy pathways include the VEGF signaling pathways; VEGF is an angiogenic cytokine that have shown to have neuroprotective properties^107^, and is implicated in metabolic syndrome^108^ and cardiovascular diseases^109^. The phosphatidylinositol pathway might be implicated in bipolar disorder and MDD, and both mood stabilizers and antidepressants have been demonstrated to act on this pathway^110^.

Another noteworthy pathway was BDNF signaling, for which numerous studies have shown its role in the pathophysiology of depression. Increased BDNF activity is also believed to be an important mechanism underlying antidepressant actions^111^. Interestingly, this pathway was listed among the top for both SSGAC-DS and MDD-CONVERGE (in the former, pathways were derived from SNPs having the same directions of risks for MDD and CV diseases while the latter involves SNPs with opposite directions of risks). However, most (except one) of the associated pathway genes were different. It is possible that the pattern of BDNF signaling differs by depression subtypes. For example, a recent study showed that BDNF levels were positively associated with the inflammatory cytokine IL-6 only in melancholic patients but not in non-melancholic subjects, with elevation of BDNF in the former group^112^. Another study on non-remitted MDD patients reported that reductions of hypersomnia, a characteristic symptom of atypical but not melancholic depression, were associated with reductions of BDNF^113^. As postulated by the authors^113^, atypical depression may be related to reduced function in the hypothalamic–pituitary–adrenal (HPA) axis, as opposed to overactivity of the HPA axis in melancholic depression^114,115^. Overactivity in the HPA system may lead to decreased BDNF expression^116^. This may explain why amelioration of symptoms in atypical depression is associated with a decrease in BDNF level. BDNF signaling may also play a role in energy balance and cardiometabolic disorders^117,118^.

A few pathways were listed among the top for MDD-CONVERGE, which were associated with reversed directions of effects for depression and CV risks. The Angiopoietin-like protein 8 regulatory pathway has been implicated in lipid metabolism and cardiometabolic disorders^119,120^. Increased serum levels of ANGPTL8 has been reported in MetS, DM and non-alcoholic liver disease^121-123^. However, this pathway has not been investigated for its role in depression. Another pathway, the oxytocin signaling pathway, plays an important role in regulating social behavior that involve the formation of attachments^124^. It has been postulated the hormone may be involved in the pathogenesis of many psychiatric disorders, including depression^125^. While studies have suggested that oxytocin may be used as a treatment for depression and anxiety ^125,126^, recent evidence showed that it may also have anxiogenic and fear-enhancing effects, enhancing the recollection of unpleasant memories^127,128^.On the other hand, oxytocin can ameliorate weight gain in diet-induced obese rats ^129,130^ and has been shown to reduce food intake in a double-blind clinical study on obese men^131^.

In summary, the shared genetic variant and pathway analyses provide insight into the molecular mechanisms underlying the comorbidities of MDD with cardiometabolic abnormalities. Given that few studies have investigated MDD comorbidities with respect to specific subtypes, the results presented here will provide a useful reference for researchers for more in-depth preclinical and clinical studies.

### Drug enrichment analyses

Drug repositioning is gaining increasingly attention in recent years as a cost-effective approach to identify novel therapies^132^. The development of every new drug requires a major investment on drug design, testing and manufacturing standards^133^. In contrast, the repositioned drugs have passed through multiple stages of clinical development, and thus have well-known safety and pharmacokinetic profiles ^134^. An example of drug repositioning in this field is pioglitazone, an insulin sensitizer which was shown in a meta-analyses of 4 RCTs to be a safe and effective adjunctive medication in non-diabetic patients with MDD^90^. To our knowledge, the current study is the first to propose and apply a human genomics approach to drug repositioning for disease *comorbidities*. The use of human genomic data for drug discovery may be particularly advantageous for psychiatric disorders, as animal models may not be able to fully mimic the human condition^135^. To model psychiatric and CV comorbidity in animals will be even more difficult.

As already discussed above, statins were ranked highly in our drug enrichment tests and some previous studies supported its therapeutic effects for both cardiovascular and depressive disorders. We will discuss several other repositioning candidates backed up by prior clinical or pre-clinical studies. Bupropion is frequently listed among the top across various depression-related phenotypes, especially when depression was considered with BMI. Interestingly, bupropion has been shown in a meta-analysis to be the only antidepressant that produces weight loss both at the acute phase and over a longer period of treatment^136^. Bupropion provides an important example supporting the validity of our approach in identifying repositioning opportunities for comorbid disorders.

Glutathione is another top repositioning hit across various depression phenotypes. It is an antioxidant and decreased level of glutathione has been reported in the prefrontal cortex of patients with psychiatric disorders, including those having MDD^137^. *N*-acetylcysteine (NAC) is a drug which increases glutathione synthesis, and it has been clinically tested and showed possible beneficial effects in a variety of neuropsychiatric disorders^138,139^. A recent meta-analysis of double-blind RCTs showed that NAC significantly improves depressive symptoms and functionality and is generally well-tolerated^140^. The benefit of glutathione or NAC in cardiometabolic diseases is less clear, but an RCT showed that NAC inhibits platelet-monocyte conjunction in type 2 DM patients, suggesting its benefit in reducing athero-thrombotic risks in these patients^141^. NAC also appeared to be efficacious in improving metabolic abnormalities in women with polycystic ovarian syndrome, with effects comparable to metformin^142^.

Several other hits are also worthy of mentioning. Allopurinol is a well-known urate-lowering agent for the treatment of gout. Interestingly, this drug has been shown to have antidepressant-like effects in mouse models^143,144^. On the other hand, studies suggested that allopurinol might also improve CV outcomes in hyperuricemic patients^145^ and in older adults with hypertension^146^. Fenofibrate is a well-known lipid-lowering agent. However, an animal study also revealed antidepressant-like effects of fenofibrate by PPAR*α* stimulation in the mesolimbic dopamine system^147^. Gemfibrozil is another repositioning hit of the fibrate class. Verapamil is a calcium channel blocker mainly used for hypertension, angina and arrhythmias. Studies have also suggested this drug might also have broader beneficial effects on cardiometabolic abnormalities. For example, verapamil has been shown to correct autophagic defects related to obesity^148^. The administration of this drug to obese mice reduced the accumulation of hepatic lipid droplets, and ameliorated pathologies of fatty liver such as inflammation and insulin resistance^148^. Regarding the effects on mood disorders, calcium channel blocker has been proposed as a novel therapeutic option for bipolar disorder and depression, although the evidence was mixed^149^. Verapamil has been reported to demonstrate antidepressant-like effects in animal models^150-152^. Interestingly, we found verapamil among the top repositioning hits for MDD-CONVERGE with fat percentage, broadly consistent with literature findings.

There are a few limitations to our study. Here we have studied GWAS datasets corresponding to four depression or depression-related phenotypes. Nevertheless, our analysis was limited to traits with available GWAS (and summary statistics); we did not cover all subtypes of depression and did not address specific symptoms of the disorder. For instance, atypical depression has not been studied via GWAS and was therefore not covered. Further GWAS or sequencing studies with detailed characterization of disease subtypes and phenotypes are warranted in view of lack of studies in this area. Also, we employed the MDD-CONVERGE sample to analyze severe melancholic depression, which is a relatively homogeneous sample comprising Chinese women^21^. Further studies are required to confirm whether our findings can be generalized to other ethnic groups or male subjects. Nevertheless, there is evidence that GWAS results from Europeans are highly replicable in East Asians and the effect sizes are also correlated^153^. In addition, as we relied on summary statistics in our polygenic score analysis, we could not easily control for covariates of interest, such as smoking. Non-linear effects are also not captured. Future studies using raw genotype data will enable more flexible analyses. Furthermore, as explained earlier, we focused on assessing genetic overlap of depression and cardiometabolic traits, but did not directly address behavioral factors (e.g. poor treatment compliance) that may be associated with the comorbidities. Further clinical studies are required to assess the magnitude and directions of effect sizes of depression phenotypes on CV risks in practice. As for the drug enrichment analyses, we have employed a relatively intuitive approach by testing for over-representation of shared genes associated with existing drugs. The significance of each gene is determined by the most significant variant and we did not take into account of LD structure and size of each gene or the significance of all SNPs in the gene. Due to the relatively moderate sample size of MDD-CONVERGE and MDD-PGC, for some traits there are no markers with tdr_11_>0.5 and hence were not included in the pathway or drug enrichment analyses. Another limitation is that we tested for over-representation of drug-related gene sets but the directions of drug effects were not explicitly considered. Delineation of the actual directions of drug actions may require more in-depth understanding of the effects of individual disease genes, and how individual drugs act on those genes. While we have presented a computational framework for drug repositioning for comorbidities, the approach should be considered exploratory rather than confirmatory, and further validation in preclinical and clinical studies are crucial before applications in the clinic.

## Conclusions

Our study highlights a significantly shared genetic basis of MDD, general depressive symptoms and neuroticism with various cardiometabolic traits. We observed positive polygenic associations with CV risks for all depression phenotypes except MDD-CONVERGE, which represents severe melancholic depression. Counter-intuitively, we observed inverse polygenic associations with CV risks for MDD-CONVERGE. Enrichment analyses of shared SNP reveal many interesting pathways, such as immune- or inflammation-related pathways, the statin pathway and BDNF signaling, that underlie the comorbidity of depressive and cardiometabolic traits. We also presented for the first time a drug repositioning approach for comorbidities using human genomics data, and many of the candidates are supported by the literature. We hope that our work will stimulate further investigations into the complex relationships between depressive and cardiometabolic diseases, and our findings will provide an important reference for future researchers.

## Acknowledgements

This work is partially supported by the Lo-Kwee Seong Biomedical Research Fund and a Direct Grant from the Chinese University of Hong Kong. We thank Professor Stephen K.W. Tsui and the Hong Kong Bioinformatics Centre for computing support.

